## Supplementary material for "Machine learning nominal max oxygen consumption from wearable reflective pulse oximetry with density functional theory": ICML ML4LMS Poster


جامعة الملك عبدالله للعلوم والتقنية
King Abdullah University of Science and Technology

### Motivation

- **Leveraging ML and wearables for precision medicine** — Recent breakthroughs in artificial intelligence have paved the way for precision medicine in unprecedented ways. The democratization and accessibility of machine learning algorithms in standard computing libraries have enabled the usage of advanced analytics to analyze large datasets that are challenging to analyze with conventional methods like regression. In this study, random forest regression (Breiman; 2001) is implemented to machine learn nominal max oxygen consumption rate, $\dot{V}O_{2,max}$ from commercial wearable sensor data based on density functional theory simulations of hemoglobin in oxygenated and deoxygenated states.
- **Inferring pulse oximetry from first principles** — Pulse oximetry is a standard physiological measure of oxygen saturation, and is typically conducted by fingertip or earlobe tests. Reflective pulse oximetry systems (Mendelson; 1983) have been developed and miniaturized to be conducted using LEDs on wearable fitness trackers, such as Fitbit, Apple Watch, WHOOP, Oura rings, and other platforms. These systems measure oxygen variation as the ratio of red to infrared ratios reflected off skin. Using density functional theory, under the independent particle assumption, Fermi's golden rule is employed to calculate absorption coefficients (Timrov; 2019) from which the absorbance ratio could be calculated from first principle simulations. Life and material science could be integrated from theory to industry with hemoglobin as an example, where the methods could be extended to other biological systems and signals from similar or other material properties from the solution to the Schrödinger many-body wave equation, posed as an Euler-Lagrange formulation of the electron density ground state energy eigenvalue problem (Schwingenschlögl; 2004).

### State-of-the-art : Random Forest

$$\hat{y}(x) = \frac{1}{n}\sum_{i=1}^{n} T_i(x) \tag{1}$$

where $\hat{y}(x)$ is the predicted value for the input $x$, $n$ is the number of trees in the forest, $T_i(x)$ is the output prediction of the leaf node reached by traversing tree $i$ using decisions based on $x$, such that:

$$T_i(x) = \begin{cases} v_{i1} & \text{if } x \text{ satisfies condition } C_{i1} \\ \vdots & \\ v_{ik} & \text{if } x \text{ satisfies condition } C_{ik} \end{cases} \tag{2}$$

where $v_{ij}$ are leaf node outputs per $i$-th tree, $C_{ij}$ are input feature conditions per node, $k$ is the number of leaf nodes in tree $i$.

### Design of First Principles Inference

- **Thm. 1:** Ground state electron density uniqueness — The ground state electron density, $\rho_0(\mathbf{r}) = N\int|\Psi_0(\mathbf{r},\mathbf{r}_2,\ldots,\mathbf{r}_N)|^2\, d\mathbf{r}_2\ldots d\mathbf{r}_N$, uniquely determines $V_{\text{ext}}(\mathbf{r})$ up to an additive constant.
- **Thm. 2:** Energy functional variational principle — $E[\rho_0] = \min_\rho E[\rho]$.
- Lead to Kohn-Sham equations by Euler-Lagrange formulation: $\left(-\frac{\hbar^2}{2m}\nabla^2 + V_{\text{eff}}(\mathbf{r})\right)\psi_i(\mathbf{r}) = \epsilon_i\psi_i(\mathbf{r})$ where $\epsilon_i$ is the Lagrange parameter approximation of the energy eigenvalue associated with the state $\psi_i$, and $V_{\text{eff}}(\mathbf{r})$ is the effective potential given by:

$$V_{\text{eff}}(\mathbf{r}) = V_{\text{ext}}(\mathbf{r}) + \int\frac{\rho(\mathbf{r}')}{|\mathbf{r}-\mathbf{r}'|}\,d\mathbf{r}' + V_{\text{xc}}(\mathbf{r}) \tag{3}$$

and $V_{\text{xc}}(\mathbf{r})$ is the functional derivative of the exchange-correlation energy $E_{\text{xc}}[\rho]$ with respect to the density, $\frac{\delta E_{xc}}{\delta\rho}$. The ground state electron density $\rho_0(\mathbf{r})$ is calculated by summing the square of the absolute value of each occupied Kohn-Sham orbital, where $N$ is the number of electrons (or occupied states):

$$\rho_0(\mathbf{r}) = \sum_{i=1}^{N}|\psi_i(\mathbf{r})|^2 \tag{4}$$

Atomic positions, $\mathbf{r}$, from structural relaxation yield inputs to calculate the complex dielectric tensor, $\epsilon(\omega)$, from which the absorption coefficient $\alpha(\lambda)$ is calculated and used to infer $\dot{V}O_{2,max}$ by learning from a random forest trained on wearable sensor data

- Stages of running a marathon:

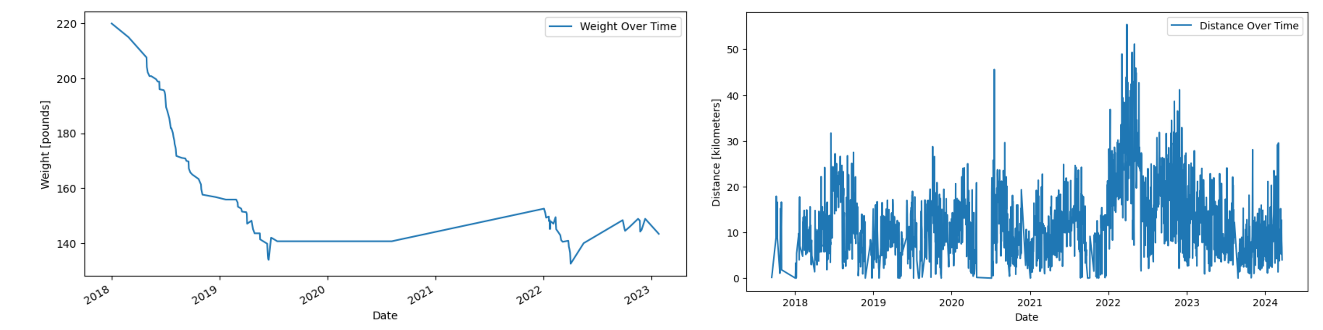


Time-series data for (top) weight loss and (bottom) distance for 5-6 years throughout marathon running in March 2022 with preparation prior and consistent running after.

- Oximetry and proprietary commercial $\dot{V}O_{2,max}$ algorithm data:

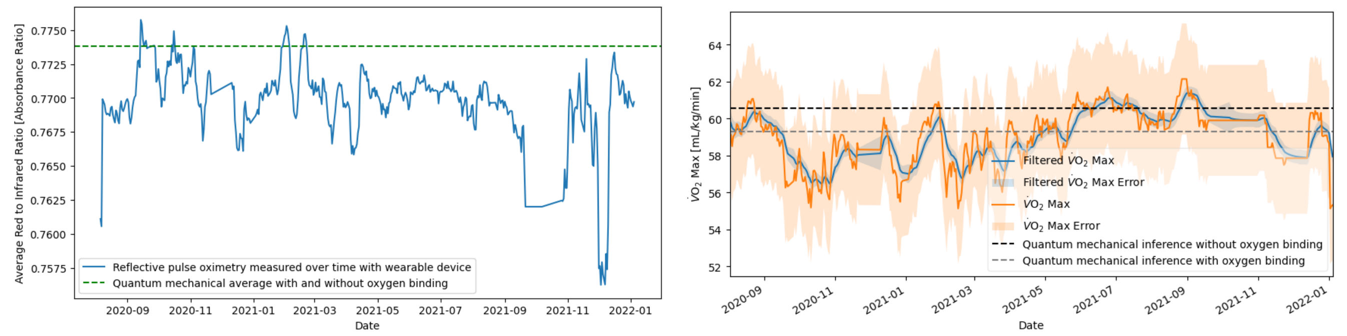

Time-series data for (left) reflective pulse oximetry and (right) max oxygen consumption rate for 1.5 years while preparing for marathon in March 2022 for nominal conditions prior.

### Learning Architecture & Hierarchy

- **Architecture:** Random forest regression model diagram

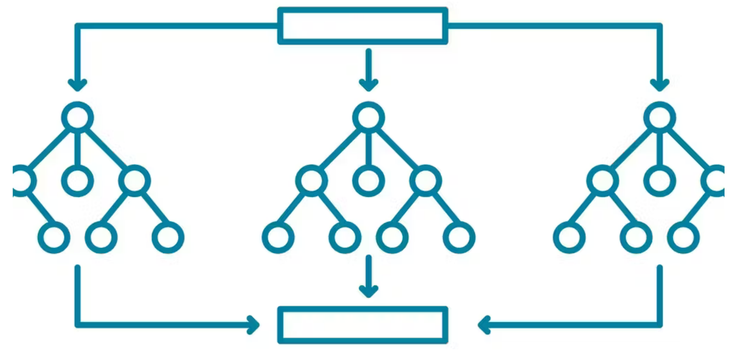

### First Principles Results

Summary of density functional theory results using the independent particle assumption-based absorption coefficient from transition state probabilities by Fermi's golden rule based on the ground state electron density dielectric tensor, $\epsilon_\rho$, where $\alpha_x$ is the $x-$nanometer absorption coefficient and $\frac{A_X}{A_Y}$ is the X-Y-nanometer absorbance ratio.

| Calculated property | Hemoglobin (Hb) | Oxyhemoglobin (HbO2) |
|---|---|---|
| $\alpha_{660nm}[\frac{1}{cm}]$ | $6.3\times10^3$ | $4.8\times10^3$ |
| $\alpha_{940nm}]\frac{1}{cm}]$ | $8.8\times10^3$ | $5.7\times10^3$ |
| $\frac{\alpha_{660nm}}{\alpha_{940nm}}$ $(\frac{A_{660nm}}{A_{940nm}})$ | 0.71 | 0.84 |

- Absorption coefficient of blood from first-principles calculations

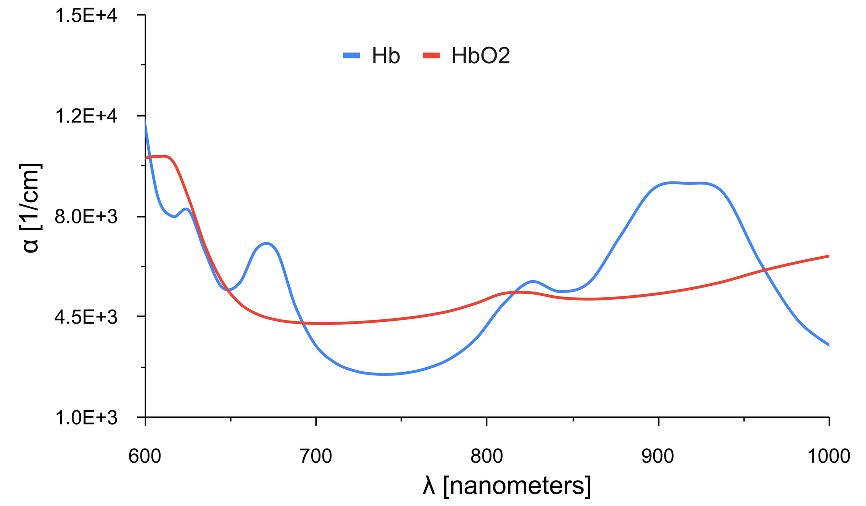


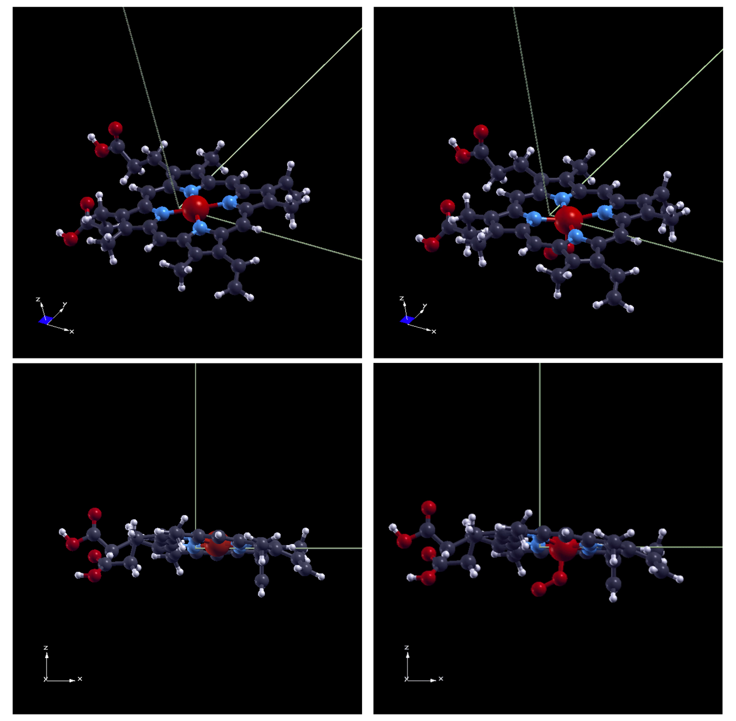

Quantum mechanical (top) absorption coefficient for the active site of human hemoglobin before (center-bottom left) and after binding oxygen (center-bottom right) at planar and side-views of the heme group.

### First Principles Inferences

Summary of machine learning results. Asterisks refers to $R^2$ for standard and reflective pulse oximetry correlation. Prediction-powered inference errors are calculated by the same models trained on the measurement errors.

| Density functional theory prediction-powered inference | Hb, A660/A940=0.71 | HbO2, A660/A940=0.84 | $R^2$ value |
|---|---|---|---|
| $SpO_2/SaO_2$ % | 96.44 | 93.15 | 0.96* |
| Linear $\dot{V}O_2$ max | $69.07 \pm 8.63$ | $47.35 \pm 12.51$ | 0.11 |
| Random forest $\dot{V}O_2$ max | $60.43 \pm 3.00$ | $58.97 \pm 3.00$ | 0.83 |
| Linear filtered $\dot{V}O_2$ max | $69.77 \pm 8.63$ | $46.65 \pm 10.50$ | 0.19 |
| Random forest filtered $\dot{V}O_2$ max | $60.71 \pm 0.82$ | $59.59 \pm 2.28$ | 0.84 |

- Learning $\dot{V}O_{2,max}$ from $\frac{\alpha_{660nm}}{\alpha_{940nm}}$ $(\frac{A_{660nm}}{A_{940nm}})$ into trained model

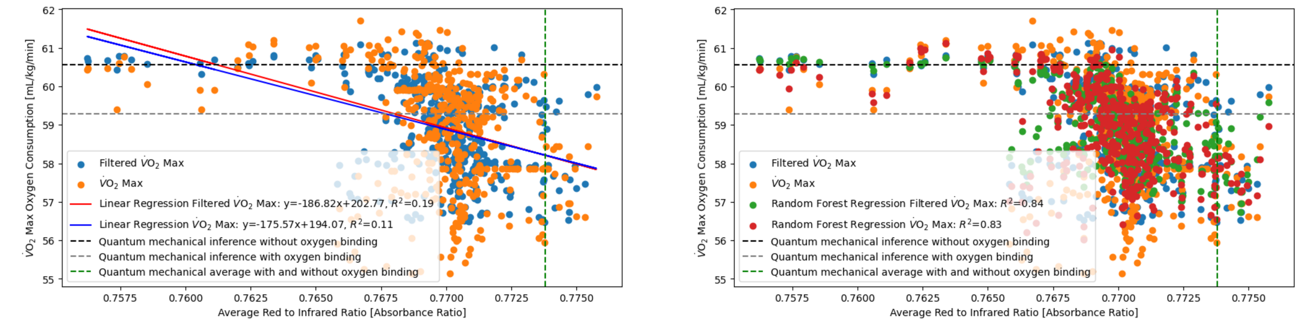

Machine learning max oxygen consumption from wearable reflective pulse oximetry with density functional theory, comparing (top) linear regression with (bottom) random forest regression to train model on wearable data for 1.5 years.

### Summary and Future Work

- This work shows that first principles calculations powered by wearable sensor data could enable AI-powered precision medicine.
- Hemoglobin is demonstrated as an example in this study and its compatibility with first principles calculation-based inferences through wearable sensor measurement models trained on physiological quantities.
- Future work: extend the approach to other biological systems and signals, such as glucose and insulin measurements.

### Software Release and References

- Code and data can be found at https://tinyurl.com/fitbit-HbO2


1 Brieman, L. (2001). Random Forests. *Machine Learning*, 45:5-32.
2 Mendelson, Y. et al. (1983). Spectrophotometric Investigation of Pulsatile Blood Flow for Transcutaneous Reflectance Oximetry. *Oxygen Transport to Tissue—IV*, pp. 93—102.
3 Schwingenschlögl, U. (2004). *The Interplay of Structural and Electronic Properties in Transition Metal Oxides*. Ph.D. Thesis, Universität Augsburg, 2004.
4 Iurii Timrov, Oscar Baseggio. Hands-on Time-Dependent Density Functional Perturbation Theory: Calculation of Absorption Spectra of Molecules. Quantum ESPRESSO Summer School, 2019. Available at: http://qe2019.ijs.si/talks/handson-day3-TDDFPT.pdf
